# Pharmacological intervention of the FGF-PTH axis as a potential therapeutic for craniofacial ciliopathies

**DOI:** 10.1101/2022.04.21.489105

**Authors:** Christian Louis Bonatto Paese, Ching-Fang Chang, Daniela Kristeková, Samantha A. Brugmann

## Abstract

Ciliopathies represent a disease class characterized by a broad range of phenotypes including polycystic kidneys and skeletal anomalies. Ciliopathic skeletal phenotypes are among the most common and most difficult to treat due to a poor understanding of the pathological mechanisms leading to disease. Using an avian model (*talpid^2^*) for a human ciliopathy with skeletal anomalies (Orofaciodigital syndrome 14), we identified disruptions in the FGF23-PTH axis that resulted in reduced calcium uptake in the developing mandible and subsequent micrognathia. While pharmacological intervention with the FDA-approved pan-FGFR inhibitor AZD4547 alone rescued expression of the FGF target *Sprouty2*, it did not significantly rescue micrognathia. In contrast, treatment with a cocktail of AZD4547 and Teriparatide acetate, a PTH agonist and FDA-approved treatment for osteoporosis, resulted in a molecular, cellular, and phenotypic rescue of ciliopathic micrognathia in *talpid^2^* mutants. Together, these data provide novel insight into pathological molecular mechanisms associated with ciliopathic skeletal phenotypes and a potential therapeutic strategy for a pleiotropic disease class with limited to no treatment options.

**Summary Statement:** Treatment options for ciliopathic phenotypes are very limited. Using an avian model, we report a novel molecular mechanism and potential therapeutic treatment for ciliopathic micrognathia.

## Introduction

Ciliopathies comprise a growing class of disorders caused by structural or functional disruptions to primary cilia (Goetz and Anderson, 2010; Plotnikova et al., 2009; Reiter and Leroux, 2017). To date, there are approximately 35 reported ciliopathies, 180 ciliopathy-associated genes, and 250 additional candidate genes (Reiter and Leroux, 2017). Ciliopathies are difficult to treat because they are pleiotropic disorders frequently manifesting in neurological, olfactory, auditory, respiratory, reproductive, excretory, and skeletal defects (Goetz and Anderson, 2010; Waters and Beales, 2011). Establishing cellular and molecular etiologies for ciliopathic phenotypes is particularly important since most ciliopathies are life-threatening diseases with limited to no treatment options (Adel Al-Lami et al., 2016).

Ciliopathic skeletal pathologies are among the most difficult of the ciliopathic phenotypes to treat for several reasons. First, these patients frequently have a very limited supply of healthy bone amenable for autograft/allograft treatment. And, even in patients with a supply of healthy bone, grafts frequently suffer from poor efficacy and substantial rejection rates (Holloway et al., 2014; Kahn, 2014). Second, current therapies geared towards inducing bone regeneration (i.e., recombinant Bone Morphogenic Protein (BMP) delivery), likely require functional cilia for signal transduction and have dangerous off-targets (Holloway et al., 2014). Finally, since very little is known regarding the cellular and molecular mechanisms that contribute to bone dysplasia in ciliopathic patients, generating pharmacological options to treat these conditions has not been possible.

One approach geared towards generating therapeutic strategies for treating ciliopathies is gaining a deeper understanding of molecular mechanisms of cilia-dependent signal transduction. The Hedgehog (Hh) pathway is perhaps the most closely linked and extensively studied pathway relative to ciliary-dependent signal transduction (Briscoe and Therond, 2013; Corbit et al., 2005; Sasai and Briscoe, 2012). Furthermore, the Hh pathway has proven to be very amenable to pharmacological intervention (Lin and Matsui, 2012; Scales and de Sauvage, 2009). Despite these promising opportunities, targeting Hh for the treatment of skeletal phenotypes is problematic due to variable Hh pathway readouts across tissues (i.e., in ciliopathies, some tissues experience a loss of Hh signaling while others experience a gain of Hh signaling) and a lack of Hh-mediated signaling during cellular processes most impacted in skeletal ciliopathies.

Several other pathways essential for skeletogenesis have been purported to utilize the cilium for signal transduction (Horner and Caspary, 2011; Kawata et al., 2021; Kunova Bosakova et al., 2019; Kunova Bosakova et al., 2018; Neugebauer et al., 2009; Wallingford and Mitchell, 2011; Yuan et al., 2019). The Fibroblast Growth Factors (FGF) pathway plays a major role in skeletogenesis, and mutations in certain ciliary proteins result in ectopic expression of genes within the FGF pathway (Kunova Bosakova et al., 2019; Kunova Bosakova et al., 2018; Mina et al., 2007; Tabler et al., 2013; Xie et al., 2020). Moreover, conditions associated with gain-of-function FGF mutations result in phenotypes reminiscent of skeletogenic ciliopathies including decreased bone mass and micrognathia (Kunova Bosakova et al., 2018; Motch Perrine et al., 2019; Zhou et al., 2013). Fgf23, a member of the endocrine subfamily of FGF ligands, is essential for bone homeostasis. Expressed in osteocytes, Fgf23 systemically interacts with parathyroid hormone (PTH) to control both bone mineralization and calcium levels throughout the body (Blau and Collins, 2015; Grau et al., 2020; Lu and Feng, 2011; Takashi et al., 2021). Misexpression of Fgf23 and PTH result in impaired bone mineralization and osteoblastic dysfunction, respectively (Iwasaki-Ishizuka et al., 2005; Lu and Feng, 2011). Interestingly, the Fgf23-PTH axis relies heavily on proper kidney function for propagation, as Fgf23 signaling induces the secretion of active Vitamin D (1,25-D3) from the kidney, which subsequently influences Ca^2+^ levels (Blau and Collins, 2015; Grau et al., 2020; Lu and Feng, 2011; Takashi et al., 2021). Although the impact of impaired FGF23-PTH signaling on bone development has been described, its correlation with skeletal phenotypes observed in ciliopathic mutants has yet to be explored.

Our previous work exploring the etiology of ciliopathic skeletal phenotypes utilized a *bona fide* avian ciliopathic model called *talpid^2^ (ta^2^*) (Abbott et al., 1959; Abbott et al., 1960). *ta^2^* embryos phenocopy the human skeletal ciliopathy Oral-facial-digital syndrome 14 (OFD14), presenting with micrognathia, hypoglossia, cleft lip/palate, hypoplastic cerebellar vermis, polydactyly, and polycystic kidneys. Genetically, just like human OFD14, *ta^2^* is caused by a mutation in the basal body protein, C2 Domain Containing 3 Centriole Elongation Regulator (C2CD3) (Chang et al., 2014). Our previous work identified impaired osteoblast maturation coupled with excessive osteoclast-mediated bone remodeling as the pathological mechanism responsible for ciliopathic micrognathia (Bonatto Paese et al., 2021). Interestingly, this mechanism is like that of osteoporosis, for which there are several pharmacological treatments.

Herein we propose a novel dual-pronged approach toward alleviating skeletal phenotypes by targeting both the molecular and cellular processes impacted during ciliopathic skeletogenesis. Our data reveal disruptions in FGF signaling, specifically within the FGF23-PTH axis in *ta^2^* embryos. This molecular profile correlates with reduced calcium uptake in the developing mandible and subsequent micrognathia. Treatment with a cocktail of AZD4547-a pan FGFR-antagonist and Teriparatide Acetate-an osteoporosis drug and PTH-agonist resulted in reduced serum Ca^2+^, increased mineralization, and increased mandibular length in *ta^2^* embryos. Together, our data suggest that a targeted approach modulating impaired FGF signaling and excessive bone degradation in ciliopathies, like OFD14, is effective in alleviating ciliopathic skeletal phenotypes.

## Results

### Ciliopathic micrognathia correlates with impaired signaling through the FGF23-PTH axis

Like several ciliopathic models, *ta^2^* embryos present with micrognathia and polycystic kidneys. Alizarin Red staining confirmed decreased bone mineralization within *ta^2^* mandibles. Transverse sections of HH39 mandibles revealed that *ta^2^* samples contained less calcium than stage-matched controls **(Fig. 1A, B).** Frontal sections through HH39 kidneys revealed several cysts within the developing *ta^2^* kidney when compared to the control **(Fig. 1C, D).** Based on the presentation of these two phenotypes, we hypothesized that the FGF23-PTH axis was impaired in *ta^2^* embryos. *FGF23* is expressed by osteocytes and osteoblasts and interacts locally with its obligatory receptor *KLOTHO* and systemically with the parathyroid hormone (*PTH*), to regulate bone mineralization and calcium metabolism. These endocrine factors induce the secretion of Vitamin D from the kidney. In normal development, vitamin D induces calcium uptake from the serum into bone **(Fig. 1E).** As per our hypothesis, impaired bone mineralization in the *ta^2^* embryos could be due to an aberrant secretion of *FGF23* and *PTH* and the polycystic phenotype could result in decreased vitamin D production, leading to decreased calcium uptake by the bone and misregulation of *FGF23* and *PTH* expression systemically **(Fig. 1E**’). To test our hypothesis, we examined the expression of genes within the FGF23/PTH axis in HH39 control and *ta^2^* kidneys and mandibles **(Fig. 1F-K).** RNAscope *in situ* hybridization showed that *KLOTHO* and *PTH* were reduced in *ta^2^* when compared to the control kidney **(Fig. 1F, G).** Concurrently, *FGF23* was significantly upregulated, and PTH was significantly downregulated when compared to control mandibles **(Fig. 1H-K).** qRT-PCR analysis confirmed *FGF23* and *PTH* were misregulated in *ta^2^* mandibles **(Fig. 1L).** The increase in *FGF23* and decrease in *PTH* expression strongly suggested aberrant calcium metabolism in *ta^2^* mutants. High-performance liquid chromatography (HPLC) for mineral contents revealed serum calcium was significantly upregulated in *ta^2^* embryos relative to controls **(Fig. 1M).** Taken together, our results revealed an imbalance in the FGF23/PTH axis which was accompanied by reduced calcium uptake in the mandible and increased calcium in the serum of *ta^2^* embryos. Based on these data, we next explored pharmacological intervention of FGF and PTH activity in *ta^2^* embryos.

**Figure 1.**
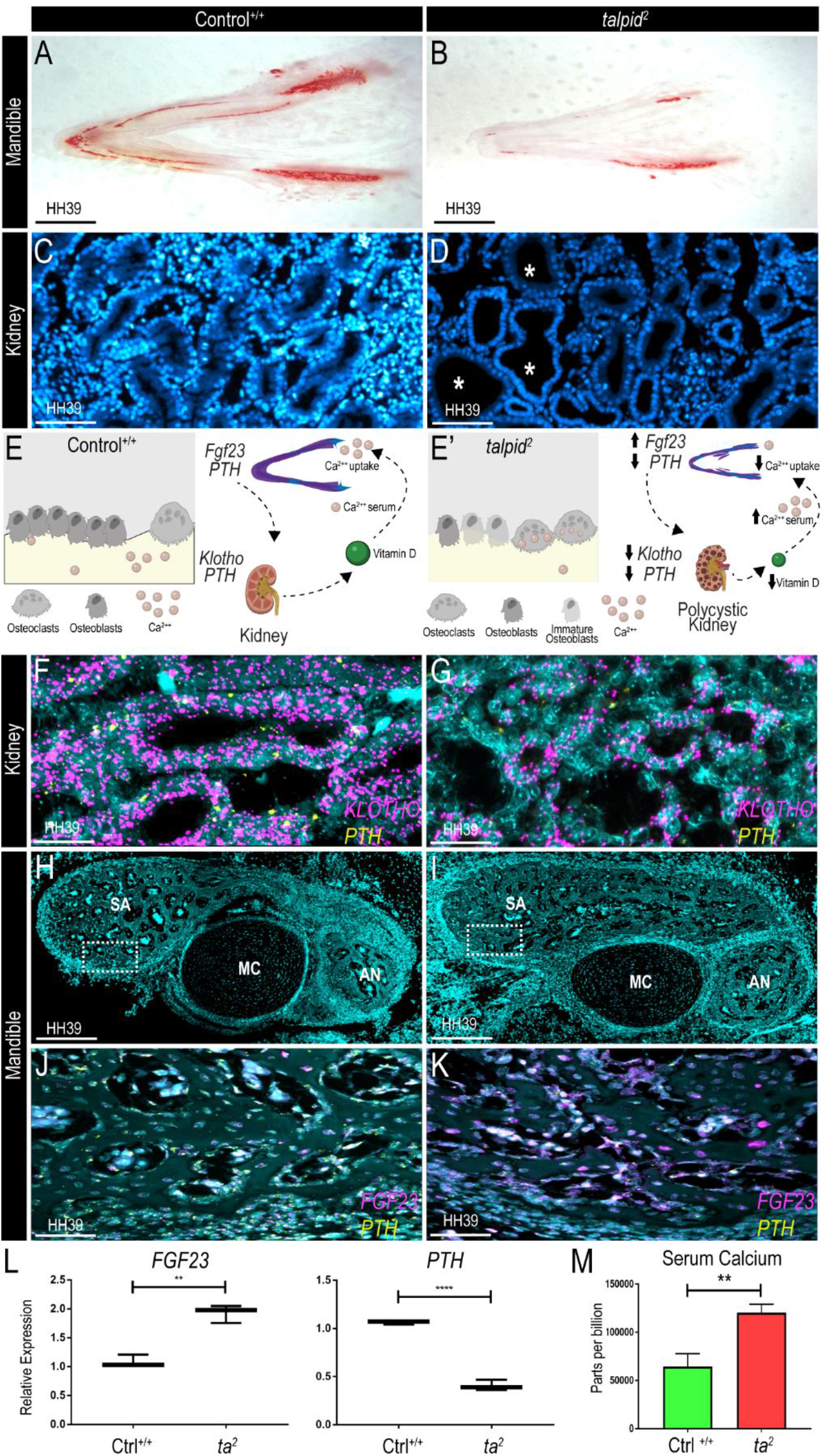
*ta^2^* mandibles and kidneys have aberrant *FGF23, KLOTHO* and *PTH* expression. (A, B) Alizarin Red stained transverse sections of HH39 control^+/+^and *ta^2^* mandibles (n=3 per group). (C, D) DAPI stained sagittal sections of HH39 control^+/+^ and *ta^2^* kidneys (white asterisks denote the presence of cystic tubules). (E) Schematic of the FGF23-PTH axis in normal embryonic development and (E’) the hypothesized axis in *ta^2^* embryos. (F, G) RNAscope *in situ* hybridization for *KLOTHO* (magenta) and *PTH* (yellow) in control^+/+^ and *ta^2^* HH39 kidney sagittal sections, nuclei counterstained for DAPI (cyan). (H, I) DAPI stained frontal sections of HH39 control^+/+^ and *ta^2^* mandible, showing the Meckel’s cartilage (MC), the angular (AN) and surangular (SA) bones. Dotted white box indicates the region of high magnification in (J, K) higher magnification pictures were taken. (J, K) RNAscope *in situ* hybridization for *FGF23* (magenta) and *PTH* (yellow) transcripts in control^+/+^ and *ta^2^* HH39 mandibular frontal sections, nuclei counterstained for DAPI (cyan). (L) qRT-PCR quantification of *FGF23* (p = 0.0016) and *PTH* (p < 0.0001) in control^+/+^ and *ta^2^* HH39 mandibles (n=3 per group). (M) Quantification of serum calcium by HPLC of control^+/+^ and *ta^2^* embryos (p = 0.0017) at HH39 (n= 3 per group). Scale bars: (A-B) 1cm (C-D) 100μm (F-G) 20μm (H-I) 100μm and (J-L) 20μm.

### Modulation of the FGF pathway alone does not alleviate ciliopathic micrognathia

FGF signaling plays a crucial role in mandibular development (Mina et al., 2007; Takashi et al., 2021; Xie et al., 2020). The master regulator of skeletal development *RUNX2* induces the expression of *FGFR2*, and this interaction is responsible for osteoblast proliferation (Kawane et al., 2018). Further, it has been shown that *FGF23* paracrine activity signals exclusively via *FGFR1*, which modulates *FGF23* expression in osteocytes (Takashi et al., 2021; Xiao et al., 2014). We evaluated the expression of *FGFR1* and *FGFR2* during osteoblast maturation (HH34) and bone remodeling (HH39). qRT-PCR revealed a significant upregulation of *FGFR2* and *FGFR1* expression and reduced expression of *SPROUTY2 (SPRY2*), a negative regulator of FGF activity, in *ta^2^* embryos at HH34 and HH39 **(Fig. 2A).** Considering these data, we attempted to rescue the micrognathic phenotype in *ta^2^* embryos by pharmacologically inhibiting FGF activity. AZD4547 is an FDA-approved, selective tyrosine kinase inhibitor that targets *FGFR1, FGFR2*, and *FGFR3* **(Fig. 2B).** To determine an effective drug dosage in HH33 embryos a dose-response curve was generated treating embryos with 10μl of either 1 or 5uM AZD4547. **(Fig. S1A-C).** Based on survival rates, 10μl of 1uM AZD4547 was utilized and delivered below the chorioallantoic membrane adjacent to the developing mandible **(Fig. 2C).** At the morphological level, we observed no significant changes between non-injected and injected *ta^2^* embryos **(Fig. 2D-G).** To determine the efficacy of AZD4547 treatment, expression of the FGF target *SPRY2* was analyzed via qRT-PCR of HH34 MNPs. Interestingly, despite a failure to rescue mandibular length, AZD4547 treatment did rescue *SPRY2* expression to that of control embryos **(Fig. 2H).**

**Figure 2.**
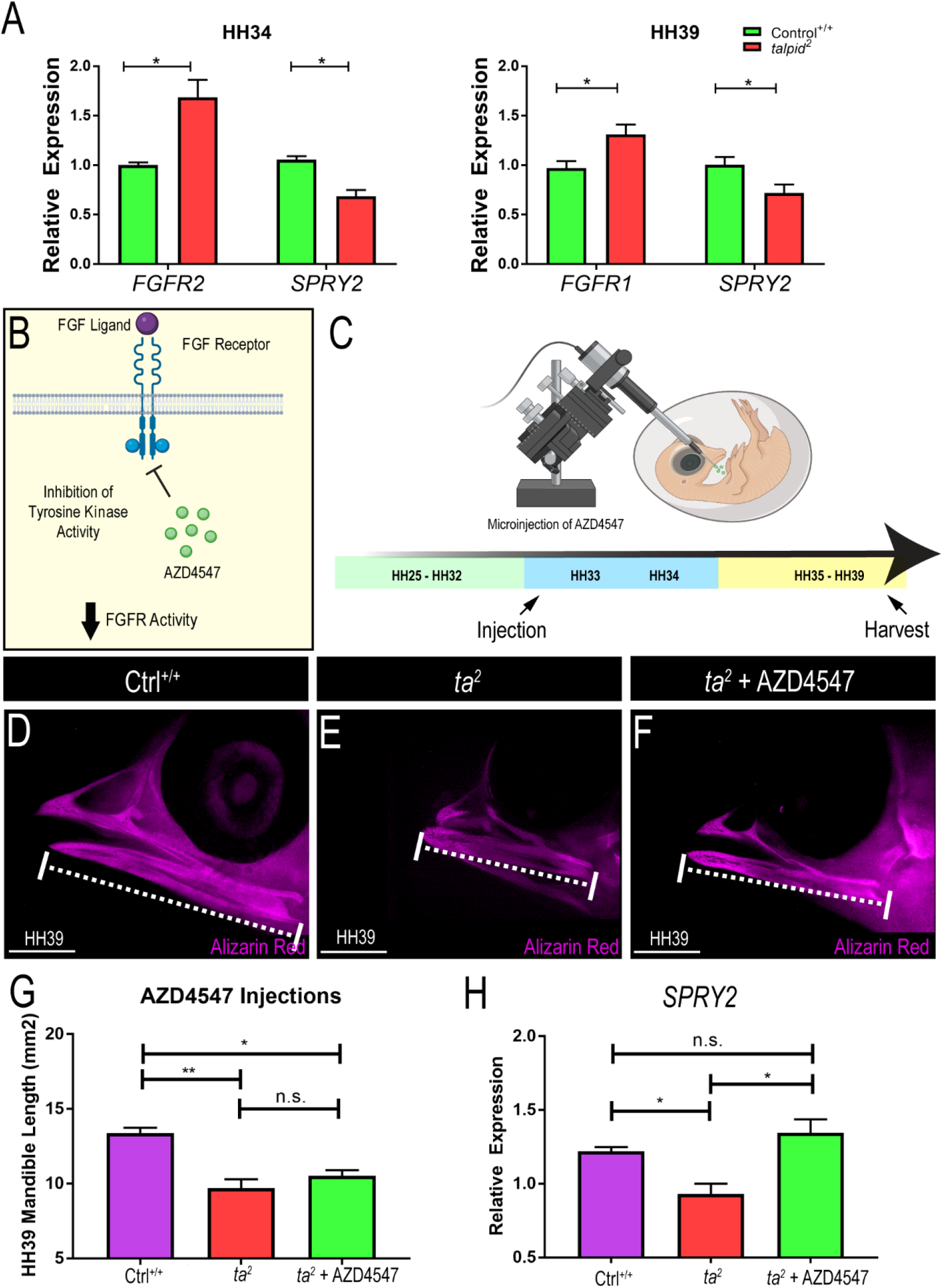
Overactive FGF signaling can be modulated with AZD4547. (A) qRT-PCR for *FGFR2* and *SPRY2* at HH34; *FGFR1* and *SPRY2* at HH39 (*p<0.05; n=4). (B) Schematic of AZD4547 mechanism. (C) Schematic of the experimental design for AZD4547 treatment. (D-F) Alizarin Red staining in HH39 ctrl^+/+^, *ta^2^* and *ta^2^* + AZD4547 treated embryos (n=4 for each group). (G) Measurements of the mandibular length of the groups depicted in D-F (*p = 0.0174; **p = 0.0084). (H) qRT-PCR quantification for *SPRY2* transcripts in the three experimental groups (*p < 0.05; n=3 per group). Data are mean±s.d. (A) Unpaired one-tailed Student’s t-test. (G-H) Ordinary one-way ANOVA. n.s, not significant. Scale bars: 2.5cm.

Our previous data revealed that increased *FGF23* expression was accompanied by decreased *PTH* expression **(Fig. 1).** PTH is crucial for the maintenance of calcium homeostasis in the body, acting directly in bone formation and resorption (Silva and Bilezikian, 2015). Thus, we next tested the potential of the PTH agonist Teriparatide Acetate to rescue ciliopathic micrognathia, using the same experimental design as previously used for AZD4547 delivery **(Fig S1A, B).** To determine an effective dosage of Teriparatide Acetate in HH33 embryos, a dose-response curve was generated treating embryos with 10μl of either 1 or 10uM of Teriparatide Acetate. Based on survival rates, 10μl of 10uM of Teriparatide Acetate was utilized and delivered as previously described **(Fig. S1B, C).** The mandibular length was not significantly increased in *ta^2^* embryos treated with Teriparatide Acetate alone **(Fig. S2C-F).** Since neither treatment alone significantly improved mandibular length, we next tested a combinatorial treatment.

### AZTeri injection is effective in alleviating ciliopathic micrognathia in *ta^2^* embryos

Given the pleiotropic nature of ciliopathies and the combinatorial cellular mechanism associated with ciliopathic micrognathia, we tested if treating *ta^2^* embryos with a cocktail of AZD4547 and Teriparatide Acetate (referred to as AZTeri from here on out) could yield a significant improvement of ciliopathic micrognathia. The AZTeri cocktail was generated using previously established dosages of individualized AZD and Teriparatide Acetate treatments (1uM). HH33 embryos were treated with 10ul AZTeri and harvested 24h later at HH34 to assess the efficacy of treatment **(Fig. 3A).** *SPRY2* expression was expanded in AZTeri treated *ta^2^* embryos, relative to untreated *ta^2^* embryos **(Fig. 3B-D).** qRT-PCR analysis validated and quantified these data and revealed that *SPRY2* expression in AZTeri treated *ta^2^* embryos was not significantly different from that observed in untreated controls **(Fig. 3E).** Western Blot analysis further revealed that AZTeri treatment was effective in downregulating MAPK cascade activity. While there was no change in total Erk levels between untreated and treated control embryos **(Fig. S3),** phospho-ERK levels were significantly downregulated in AZTeri treated *ta^2^* embryos when compared to the untreated *ta^2^* group **(Fig. 3F).**

**Figure 3.**
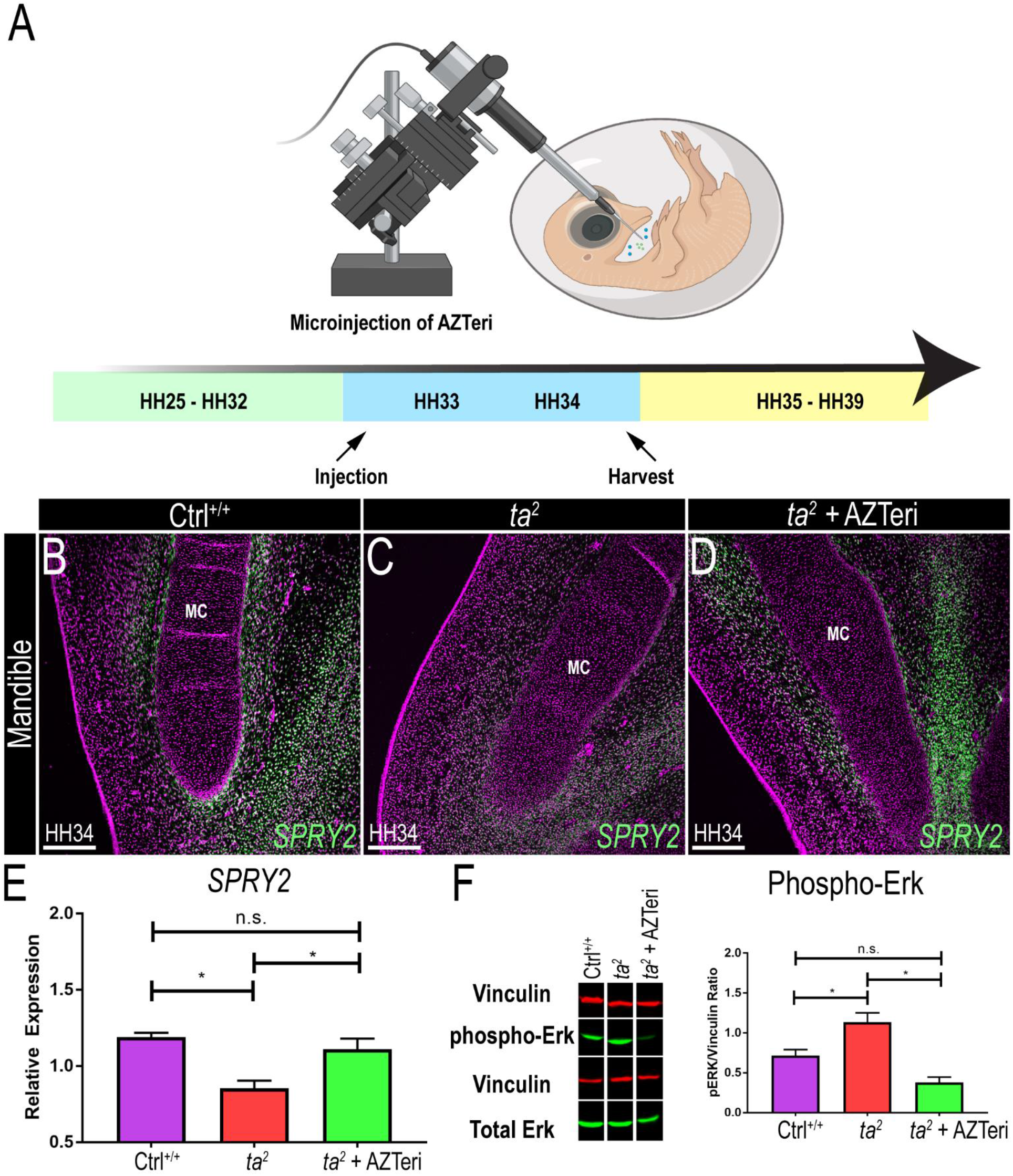
AZTeri treatment in the *ta^2^* mandible. (A) Schematic of the experimental design for AZTeri treatment. (B-D) RNAscope *in situ* hybridization for *SPRY2* (green) in ctrl^+/+^, *ta^2^* and *ta^2^* + AZTeri transverse mandibular sections (n=4 per group). (E) qRT-PCR quantification for *SPRY2* transcripts in the three experimental groups (n=4 per group). (F) Western blot for phosphorylated-ERK and total ERK, and quantification of pERK/Vinculin ratio in non-injected ctrl^+/+^, *ta^2^* and *ta^2^* + AZTeri embryos at HH34 (n=3 per group). Nuclei counterstained for DAPI (magenta). MC: Meckel’s cartilage. Data are mean±s.d. *P<0.05 (Ordinary one-way ANOVA). n.s, not significant. Scale bars: 200μm.

To test the potential of AZTeri as a therapeutic agent for skeletal ciliopathies, HH33 embryos were treated with 10μl of AZTeri and harvested at HH39. AZTeri-treated *ta^2^* embryos demonstrated a significant increase in mandibular length when compared to untreated *ta^2^* embryos **(Fig. 4A-D).** Transverse sections through HH39 mandibles revealed a robust amount of calcium incorporated into the mandible of controls relative to *ta^2^* mandibles **(Fig. 4E, F).** Interestingly, AZTeri treated *ta^2^* embryos showed a marked improvement in the amount of mandibular calcium pockets compared to the untreated *ta^2^* sections **(Fig. 4G).** Measurements of the calcified area demonstrated that AZTeri treatment restored the amount of calcified tissues in *ta^2^* to that of control embryos **(Fig. 4H).** AZTeri-treated *ta^2^* embryos also had reduced bone remodeling, as assayed by tartrate-resistant acid phosphatase (TRAP) staining, when compared to the untreated *ta^2^* embryos **(Fig. 4I-L).** Moreover, HPLC analysis of AZTeri-treated *ta^2^* embryos further revealed that the increase of calcium and decrease of TRAP staining in the developing mandible correlated with decreased serum calcium levels **(Fig. 4I).** Finally, to confirm if PTH levels and osteoclast activity were indeed changed in the treated embryos, we performed RNAscope in situ hybridization in frontal sections of HH39 mandibles for *PTH* and *SPP1* transcripts. *PTH* expression was reduced, while *SPP1* expression was increased in *ta^2^* mandibles, relative to controls. AZTeri treatment resulted in increased *PTH* expression and reduced *SPP1* expression when compared to untreated *ta^2^* embryos **(Fig. 4N-S),** further suggesting that excessive bone remodeling is partially alleviated in treated embryos **(Fig. 4Q-S).** Taken together, these results demonstrated the potential of AZTeri treatment for ciliopathic micrognathia.

**Figure 4.**
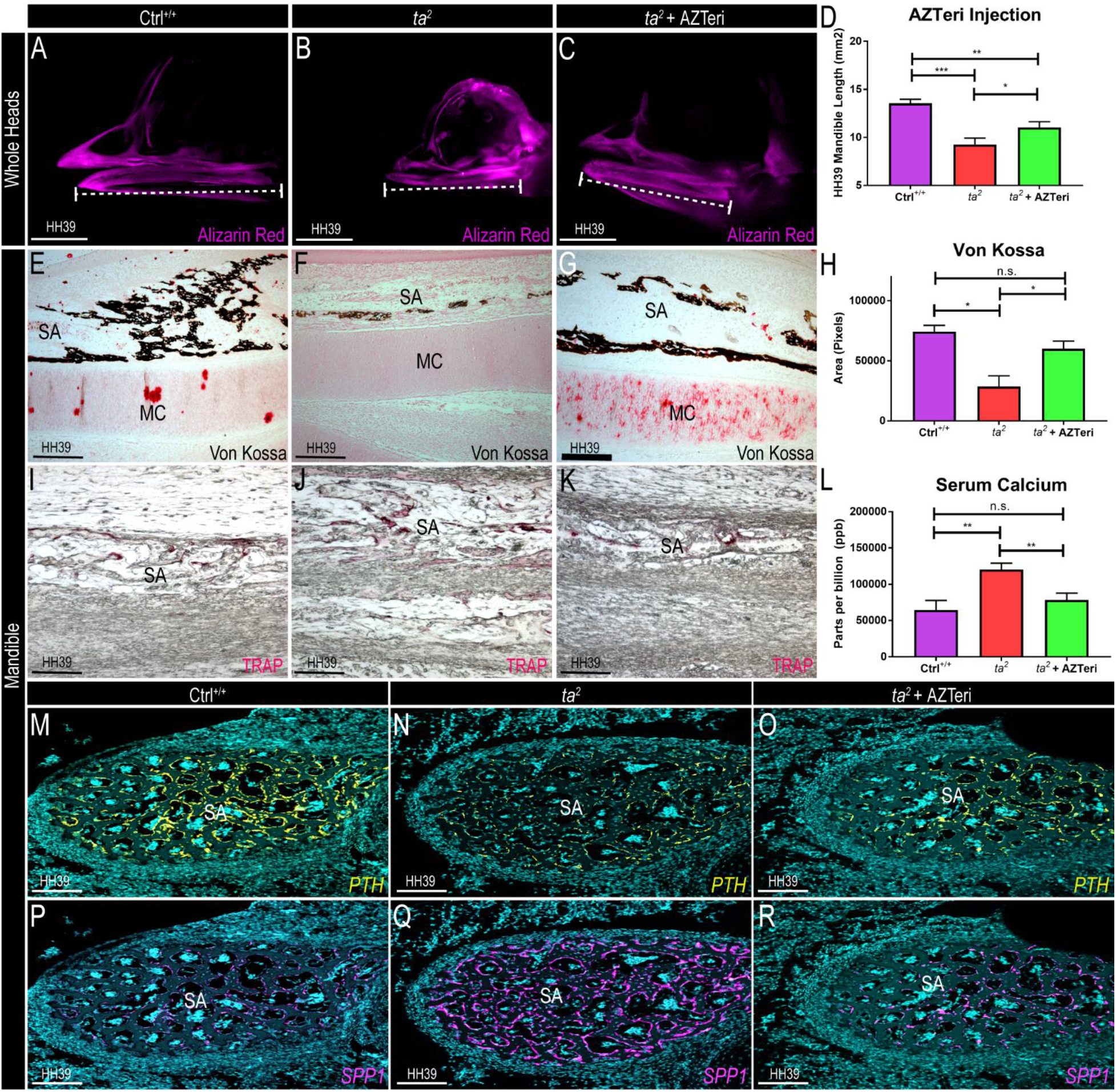
AZTeri treatment alleviates the micrognathic phenotype in *ta^2^* embryos. (A-C) Alizarin Red stained heads at HH39 of ctrl^+/+^, *ta^2^* and *ta^2^* + AZTeri embryos (n=3 for each group). (D) Measurements of the mandibular length of ctrl^+/+^, *ta^2^* and *ta^2^*+ AZTeri embryos (*p= 0.0199; **p= 0.0040; ***p= 0.0002). (E-G) Von Kossa staining in transverse sections of ctrl^+/+^, *ta^2^* and *ta^2^ +* AZTeri HH39 mandibles (n= 4 per group). (H) Area quantification of Von Kossa stained HH39 mandibular sections (*p< 0.05). (I-K) TRAP staining in transverse sections of ctrl^+/+^, *ta^2^* and *ta^2^* + AZTeri HH39 mandibles (n= 4 per group). (L) Quantification of serum calcium by HPLC of ctrl^+/+^, *ta^2^* and *ta^2^* + AZTeri HH39 embryos (**p< 0.05; n=3 per group). (M-O) RNAscope *in situ* hybridization for *PTH* (yellow) in ctrl^+/+^, *ta^2^* and *ta^2^* + AZTeri HH39 mandibular frontal sections (n=3 per group). (P-R) RNAscope *in situ* hybridization for *SPP1* transcripts (magenta) in ctrl^+/+^, *ta^2^* and *ta^2^* + AZTeri HH39 mandibular frontal sections (n=3 per group). Data are mean±s.d. (ordinary one-way ANOVA). n.s, not significant. Scale bars: (A-C) 2.5cm, (E-G) 200μm, (I-L) 50μm and (N-S) 20μm.

## Discussion

Herein, we present a potential avenue for the pharmacological intervention of ciliopathic skeletal phenotypes. Utilizing the *ta^2^* avian mutant as a model for a human ciliopathy, we identified disruptions in the FGF23/PTH signaling axis concomitant with decreased bone mineralization and increased serum calcium. These data, in concert with our previous reports that excessive bone resorption contributed to ciliopathic micrognathia (Bonatto Paese et al., 2021), informed our hypothesis that treatment simultaneously targeting FGF signaling and bone resorption would rescue micrognathia in *ta^2^* embryos. These findings support a potential drug-based therapeutic option for human ciliopathy patients.

Avians are an exquisite model for pharmacological testing due to *in ovo* embryonic accessibly, low cost, and an abundant number of embryos (Zosen et al., 2021). Several drugs currently used in preclinical cancer trials or treatments were initially tested on avian embryos (Bueker and Platner, 1956; Karnofsky and Lacon, 1964; Kue et al., 2015; Ryley, 1968; Zuniga et al., 2003). While in *ovo* screens have provided a wealth of information on toxicity and off-target effects, the lack of avian models for human disease has prevented more robust usage of the egg as a tool for testing pharmacological agents in human health research.

The *talpid^2^* is perfectly suited for such studies. First, it phenocopies human ciliopathies on both a genetic and biochemical level and survives well into development. Second, since most ciliopathic models are early embryonic lethal, murine conditional knock-out models are commonly used to study molecular mechanisms. While this is effective in examining a ciliopathic insult on one tissue, it fails to consider the pleiotropic nature of ciliopathies as they present in human patients. As such, the *talpid^2^* represents a unique and powerful model that is not only easily accessible but also highly representative of a human ciliopathy (Bonatto Paese et al., 2021; Chang et al., 2014; Schock et al., 2015).

One of the most common skeletal phenotypes associated with ciliopathies is micrognathia. Micrognathia significantly impacts a patient’s ability to breathe, eat and speak. Treatment options for micrognathia are limited. Surgical procedures, like distraction osteogenesis, are highly invasive and the poor quality of the bone in ciliopathy patients makes treatment like this less effective (Abramson et al., 2013; Breik et al., 2016; Holloway et al., 2014; Kahn, 2014; Perlyn et al., 2002; Tomonari et al., 2017). To eliminate the need for surgical intervention, pharmacological treatments for micrognathia have been explored. Drug treatments for osteoporosis were seen as strong candidates for treatment. Osteoporosis is broadly defined as “an imbalance between bone formation and bone resorption” (Bodenner et al., 2007). Mechanistically, this description is very similar to the pathology observed in the *ta^2^* mandibles (Bonatto Paese et al., 2021). Bisphosphonates represent potent inhibitors of bone resorption that are FDA approved for the treatment of osteoporosis. In an avian model, bisphosphonate treatment significantly elongated the mandible (Ealba et al., 2015). Despite the efficacy of bisphosphonate treatment in avians, treatment in humans has proven less effective and has been associated with the development of bisphosphonate-related osteonecrosis of the jaw (BRONJ) (Eckert et al., 2007; Rayman et al., 2009). Thus, additional experiments focusing on alternative pharmacological treatments for micrognathia are necessary.

Teriparatide Acetate (TA), a component of the AZTeri treatment used herein, represents another FDA-approved treatment for osteoporosis. TA effectively reduces bone resorption and has shown promising results in phase 4 trials (Leder, 2017). TA has been successfully used for the treatment of BRONJ (Chopra and Malhan, 2020; Dos Santos Ferreira et al., 2021; Kwon and Kim, 2016; Sim et al., 2020; Yu and Su, 2020), and reduced serum calcium levels and improved bone integrity in osteoporosis and hypoparathyroidism patients (Gutierrez-Cerecedo et al., 2016; Satterwhite et al., 2010). Considering the variable efficacy and side effects in human patients, it will be important to carefully examine other osteoporosis-approved drugs (Denosumab, etc.) for the treatment of ciliopathic skeletal phenotypes (Tsai et al., 2019).

In addition to targeting the cellular process of bone resorption with TA, we also hypothesized that treating excessive FGF activity would prove necessary for the treatment of micrognathia. Previous results revealed an association between ciliopathies and FGF syndromes, however; the association was specifically between FGF signaling and the onset of maxillary phenotypes, like high arched palate (Tabler et al., 2013). Mandibular ciliopathic phenotypes, on the other hand, have been more commonly associated with aberrant Hh or Wnt signaling (Elliott et al., 2018; Millington et al., 2017; Zhang et al., 2011). While much of the data on FGF and mandibular development focuses on an early patterning role of FGF8 (Mina et al., 2007; Shigetani et al., 2000; Terao et al., 2011; Zhou et al., 2013), FGF23 plays an important role later in skeletal development by modulating parathyroid hormone and calcium signaling (Blau and Collins, 2015; Lu and Feng, 2011). As Hh and Wnt signaling have numerous roles throughout the embryo at this stage of skeletogenesis, focusing specifically on FGF23 signaling may prove to be the most targeted mode of treatment for pleiotropic diseases, like ciliopathies, with skeletal phenotypes.

Calcium signaling plays a pivotal role during bone development, and depleted calcium uptake is the main cause of conditions such as osteoporosis and rickets (Monsen, 1989). There is no consensus as to if the primary cilium plays a major role in calcium signaling (Delaine-Smith et al., 2014; Delling et al., 2013; Delling et al., 2016; Hoey et al., 2012; Lee et al., 2015; Malone et al., 2007; Saternos et al., 2020), yet our results support a systemic role for cilia in the differentiation of osteoblasts (Bonatto Paese et al., 2021). It is possible that the role of cilia in calcium uptake may vary between tissues (e.g., node vs. osteoblast), temporally during development, or between chemosensory and mechanosensory cilia. More detailed experiments will need to be done to definitively determine the relationship between the cilium and calcium uptake in the developing mandible.

In summary, our work proposes a novel molecular mechanism and treatment strategy for ciliopathic micrognathia using a cocktail of FDA-approved drugs. While this treatment does not completely restore mandibular length to that of control embryos, it does significantly rescue the micrognathic phenotype. As a complete rescue of micrognathia may be optimistic at this time, a realistic goal for this treatment option is to restore the mandible to a length that alleviates the need for repeated, invasive surgeries and allows patients a better quality of life.

## Material and Methods

### Embryonic collection and genotyping

Fertilized control and *ta^2^* eggs were purchased from the University of California, Davis. Eggs were incubated at 38.8°C in a rocking incubator with humidity control. Staging followed the Hamburger-Hamilton staging system, and genotyping was performed as previously described (Bonatto Paese et al., 2021; Hamburger and Hamilton, 1951). Unless noted otherwise in the figure legend, every experiment utilized five embryos for each experimental group.

### Skeletal staining

Samples were incubated in 0.005% Alizarin Red S (Sigma-Aldrich A5533) in 1% KOH for 3 hours at room temperature and cleared in 1% KOH. Once cleared, samples were incubated in Glycerol:KOH 1% (50:50) solution. For imaging and long-term storage, samples were kept in 100% glycerol. Stained specimens were imaged using a Leica M165 FC stereo microscope system.

### qRT-PCR

RNA was extracted using TRIzol reagent (Invitrogen) and cDNA was synthesized using SuperScript III (Invitrogen). HH39 mandibles were first frozen with liquid nitrogen and ground using a mortar and pestle to ensure homogenous extraction. SYBR Green Supermix (Bio-Rad) and a Quant6 Applied Biosytems qPCR machine were used to perform qRT-PCR. All the genes were normalized to GAPDH expression. Negative controls were performed by omitting the cDNA in the mixture. The level of expression for each gene was calculated using the 2^-ΔΔCq^ method (Livak and Schmittgen, 2001). Unpaired one-tailed Student’s t-test was used for statistical analysis. P<0.05 was determined to be significant.

### RNAScope *in situ* hybridization

RNAscope in situ hybridization was carried out as previously described (Bonatto Paese et al., 2021). The transcripts used in this study were as follow: *FGF23* (ACD - 1002831), PTH (ACD - 1003861), *SPP1* (ACD - 571601) and *SPRY2* (ACD - 1086991), were detected using the RNAscope Multiplex Fluorescent V2 kit per manufacturer’s instructions. Both sections and wholemount samples were imaged using a Nikon A1 LUN-V inverted microscope system.

### Embryonic treatment

Three mixes were utilized in this study: AZD4547 (Selleck Chem - S2801) was diluted to 1μM in 4% DMSO+30% PEG 300+5% Tween 80+ddH2O. Teriparatide Acetate (Selleck Chem - P1033) was diluted to 1μM in ddH2O. AZTeri was a mix of 1μM of AZD4547 and 1μM of Teriparatide Acetate diluted in ddH2O. Embryos were treated at HH33 embryos via applying 10μl of the drugs under the chorioallantoic membrane immediately adjacent to the mandible. Embryos were then incubated without shaking in the incubator. Wholemount heads were dissected at either HH34 or HH39 and processed for further analysis.

### Analysis of serum calcium content

100μl of blood was collected from the vitelline vein of HH39 embryos was collected on ice with microcapillaries, weighted and sent for processing by the R. Marshall Wilson Mass Spectrometry facility at University of Cincinnati. Inductively Coupled Plasma - Mass Spectrometry with High Performance Liquid Chromatography (ICP-MS HPLC) was utilized.

### Histological analysis

Hematoxylin and eosin (H&E) staining was performed using standard protocols. For calcium deposit analysis, 7μm transverse sections of HH39 mandibles were used with the Von Kossa Stain Kit (Calcium Stain) (Abcam - ab150687), following manufacturer instructions. TRAP staining was performed on 8 μm thick transversal sections of undecalcified HH39 mandibles using the Acid Phosphatase Leukocyte (TRAP) kit (Sigma-Aldrich, 387A) following the manufacturer’s protocol.

### Western Blot

Embryos were injected at HH33, and mandibles were dissected at HH34 for processing. Collected tissue was sonicated in cold RIPA buffer (50 mM Tris-HCl, pH 7.4, 1% NP-40, 0.25% sodium deoxycholate, 150 mM NaCl, 1 mM EDTA) containing protease and phosphatase inhibitors (ThermoFisher 78440). The protein extract was collected after 10 minutes full-speed centrifugation at 4C. 20ug of protein from each embryo was used for Western blot, with the following primary and secondary antibodies: ERK1/2 (Cell Signaling Technology - 9101S, 1:1000), Phospho-p44/42 MAPK (ERK1/2) (Novus Biologicals NB110-96887, 1:1000), Vinculin (Santa Cruz Biotechnology sc-73614, 1:2000), IRDye^®^ 800CW Donkey anti-Rabbit IgG (LICOR 926-32213, 1:2000), IRDye^®^ 680RD Donkey anti-Mouse IgG (LICOR 925-68072, 1:2000). Images were taken by LICOR Odyssy^®^ DLx. Densitometry was done by ImageJ.

### Statistical methods

Unpaired t-tests (two groups) or one-way ANOVA (three and four groups) were used in comparisons for statistical analysis between groups. It was considered significant when the two-tailed analysis were p<0.05.

## Acknowledgements

We would like to thank the University of California – Davis avian facility (Mary Delany, Jackie Pisenti and Kevin Bellido) for maintenance and husbandry of the *talpid^2^* colony. Technical assistance was given by Dr Matt Kofron for image acquisition and analysis (Confocal Imaging Core at Cincinnati Children’s Hospital Medical Center) and Dr Julio Landero Figueroa for calcium serum analysis (University of Cincinnati - R. Marshall Wilson Mass Spectrometry facility). We are grateful for the Brugmann laboratory technical assistance and feedback. This study was funded by the National Institute of Dental and Craniofacial Research (R35 DE027557) and to S.A.B. and the Cincinnati Children’s Hospital Medical Center internal grant for C.L.B.P. (Arnold W. Strauss Fellowship).

## Competing interests

The authors declare no competing or financial interests.

## Author contributions

Conceptualization: C.L.B.P and S.A.B.; Methodology: C.L.B.P., D. K., and C.F.C.; Validation: C.L.B.P., C.F.C.; Formal analysis: C.L.B.P. and S.A.B.; Investigation: C.L.B.P., C.F.C., Resources: S.A.B.; Writing - original draft: C.L.B.P., C.F.C and S.A.B.; Writing - review & editing: C.L.B.P., C.F.C., S.A.B.; Visualization: C.L.B.P., D.K., C.F.C., S.A.B.; Supervision: S.A.B.; Project administration: S.A.B.; Funding acquisition: C.L.B.P. and S.A.B.

---

**Supplementary Figure 1.** (A) Dose response curve of AZD4547 treatment. X-axis represents drug concentration (uM) and Y-axis represents the mortality rate (%) of total embryos treated. (B) Dose response curve of Teriparatide Acetate treatment. X-axis represents the drug concentration (uM) and Y-axis represents the mortality rate (%) of total embryos treated. Yellow-dashed line shows the 50% mortality rate, and the red-dashed line represents the chosen concentration. (C) Table containing the number of embryos treated for each drug concentration of AZD4547 and Teriparatide Acetate.

**Supplementary Figure 2.** (A) Schematic of the mechanism of action for Teriparatide Acetate. (B) Experimental design for Teriparatide Acetate treatment. (C-E) HH39 Alizarin Red stained heads of ctrl^+/+^, *ta^2^* and *ta^2^* + Teriparatide Acetate embryos (n=3 for each group). (F) Measurements of the mandibular length of the groups depicted in C-E (p> 0.05). Data are mean±s.d. (ordinary one-way ANOVA). n.s, not significant. Scale bars: 2.5cm (C-E).

**Supplementary Figure 3.** (A) Western blot for phosphorilated-ERK and total ERK, and quantification of pERK/Vinculin ratio in ctrl^+/+^ and ctrl^+/+^ + AZTeri embryos at HH34 (n=3 per group). Data are mean±s.d. *P<0.05 (unpaired one-tailed Student’s t-test).

## Notes

### Competing Interest Statement

The authors have declared no competing interest.

